# Type VII secretion system and its effect on Group B Streptococcus virulence

**DOI:** 10.1101/2023.01.23.525132

**Authors:** Yulia Schindler, Galia Rahav, Israel Nissan, Gal Valenci, Miriam Ravins, Emanuel Hanski, Dana Ment, Dorit Tekes-Manova, Yasmin Maor

## Abstract

GBS may cause a devasting disease in newborns. In early onset disease of the newborn the bacteria are acquired from the colonized mother during delivery. We characterized type VII secretion system (T7SS), exporting small proteins of the WXG100 superfamily, in group B Streptococci (GBS) isolates from pregnant colonized women and newborns with early onset disease (EOD) to understand better understand T7SS contribution to virulence in these different clinical scenarios.

GBS isolates were obtained from colonized mother prior to delivery and from newborns with EOD. DNA was analyzed for T7SS genes. A mutant EOD strain (ST17) was created by knocking out the *essC* gene encoding a T7SS protein. *Galleria mellonella* larvae were used to compare virulence of colonizing, EOD, and mutant EOD isolates.

33 GBS genomes were tested, 17 EOD isolates and 16 colonizing isolates. The T7SS locus encoded 8 genes: *essC*, membrane-embedded proteins (*essA; essB*), modulators of T7SS activity *(esaA; esaB; esaC*) and effectors: [*esxA* (SAG1039); *esxB* (SAG1030). ST17 isolates encode two copies of the *essC* gene and *esxA* gene encoding putative effectors but were present only in 23.5% of isolates. In ST1 isolates three copies of esxA gene were identified, but in ST6 and ST19 isolates all T7SS genes were missing. EOD isolates demonstrated enhanced virulence in *G. mellonella* model compared to colonizing isolates. The 118659Δ*essC* strain was attenuated in its killing ability, and the larvae were more effective in eradicating 118659Δ*essC* infection. *essC* gene deletion was associated with reduced bacterial growth. We demonstrated that T7SS plays an essential role during infection and contributes to GBS pathogenicity.

**Author Summary:** Type VII secretion system (T7SS) is related to virulence in various bacteria but is not well characterized in Group B Streptococci (GBS). GBS may cause sepsis, meningitis, and death in newborns. The bacteria rarely cause disease in pregnant mothers. Newborns acquire GBS from the colonized mother during delivery. We studied the role of T7SS in GBS isolates obtained from newborns with GBS sepsis in the first week of life and in colonized pregnant mothers. By studying T7SS genes we discovered that the genetic structure of the T7SS differs between isolates causing severe disease and colonizing isolates. To study the virulence of different GBS isolates we injected them into larvae and monitored larvae survival. Isolates causing severe disease in the newborn caused a more severe disease in larvae compared to colonizing isolates. We then deleted T7SS genes in GBS isolates causing severe disease. The killing activity of GBS isolates without T7SS genes was attenuated. The larva responded to these bacteria similarly to the response found when injecting the larva with GBS isolates from colonized mothers. These results support our hypothesis that T7SS is important for causing severe infection in the newborn and that this system contributes to GBS pathogenicity.

## INTRODUCTION

Group B streptococcus (GBS) also known as *Streptococcus agalactiae* is a commensal bacterium that belongs to the human microbiota colonizing the gastrointestinal and genitourinary tract^1^. In most cases the colonization in humans is harmless but GBS can also cause severe disease^2,3^. An important manifestation of GBS disease is neonatal sepsis and meningitis^3^. Early-onset disease (EOD) in the newborn is a devastating disease that results from the vertical transmission of GBS from colonized mothers through contaminated amniotic or vaginal secretions to her newborn. GBS isolates can be divided into10 distinct serotypes (Ia; Ib; II-IX) based on a serological reaction directed against the polysaccharide capsule^1^. Based on multilocus sequence typing (MLST) most human GBS isolates can be clustered into six major sequence types (STs)^1^.

GBS has a variety of putative virulence factors that facilitate its ability to cause disease, some of which have been identified and characterized^4,5^.

Bacterial pathogens utilize a multitude of methods to invade mammalian hosts, damage tissue sites, and escape the immune system^6^. One essential component for many bacterial pathogens is secretion of proteins across phospholipid membranes^7^. Type VII secretion system (T7SS) is a specialized secretion system in Gram positive bacteria first discovered in *Mycobacterium spp*, where it is responsible for the export of small proteins that are members of the WXG100 superfamily^8^. In *Mycobacterium tuberculosis* T7SS plays an important role in bacterial virulence and persistence of infection^9–11^. Analogous substrates and some components of these systems have also been identified in several other Gram-positive organisms, including *Staphylococcus aureus, Streptococcus pyogenes, Streptococcus pneumoniae and Bacillus anthracis*^12–14^.

There are commonalities and differences between the T7SS of *Actinobacteria* and *Firmicutes*^15,16^. A membrane-embedded ATPase of the FtsK/SpolllE family termed EssC is found in all T7SSs. In both systems the protein shares a similar overall topology, with two transmembrane domains that are usually followed by three P-loop ATPase domains at the C-terminus, that energize substrate secretion. The ATPase domain of EssC interacts with conserved WXG100 protein substrates, through a signal sequence^17^. The second common component is at least one small protein of the WXG100 family, EsxA, which is secreted by the T7SS ^18^. In Mycobacteria, EsxA homologues are secreted as heterodimers with EsxB (LXG-domain containing protein)^19,20^, whereas in Firmicutes EsxA is secreted as a homodimer^21,22^. The T7SS is encoded by the ess locus. In addition to EsxA and EssC, further integral and peripheral membrane proteins encoded by the locus (such as EsaA, EssA, EssB and EsaB). In *S. aureus* they are also essential components of the secretion machinery^12,23^. Additionally, increasing numbers of reports have shown a role for the T7SS and/or EsxA in the pathogenesis of several Gram-positive bacteria^18,24,25^; however, there is insufficient data regarding the structure and distribution of T7SS in clinical GBS strains. Recently the structure of T7SS in GBS strains was characterized and four T7SS subtypes based on the C-terminus of the ATPase EssC were identified^26^. Additionally, the genetic diversity of the T7SS in GBS isolates was also identified^27^, but the clinical significance of this secretion system in GBS is still unknown.

We recently demonstrated that in a population of orthodox Jews treated at Maayaney Hayeshua Medical Center (MHMC) serotype III [sequence type (ST)17] was the most common serotype in EOD while serotype VI (ST1) was the prevalent serotype among colonizing isolates^28^. This prompted us to search for the presence and structure of the T7SS locus among these clinical GBS isolates and to assess the effect of the T7SS on the virulence of isolates causing colonization and isolates causing invasive disease (EOD).

## RESULTS

Thirty-three GBS isolates were studied, 17 from neonates with EOD and 16 from asymptomatic pregnant women. ST types were: ST17 (n=17) from neonates with EOD, ST1 (n=12), ST19 (n=3) and ST6 (n=1) from asymptomatic pregnant women.

### Identification of three GBS T7SS subtypes among various ST’s

We analyzed the structure of the T7SS locus compared to the reference genome *S. agalactiae* 2603 V/R. We observed structures related to T7SS in all isolates except in ST19 and ST6 isolates. Furthermore, we found an extensive amount of genetic diversity in T7SS operons regarding sequence homology of core genes and putative effectors genes (Table 1). T7SS core genes *EsaA, EssA, EssB*, and EsaB were found homologous to those found in *S. aureus* genomes^37^ and had >96% identity to the reference strain. The difference between the isolates was in the sequence homology of *essC* genes (encodes FtsK/SpoIIIE-type ATPase) and the presence of one or more *esxA* homologs, and the presence of putative LXG toxin/anti-toxin-encoding gene *(esxB*). Based on these results we suggested three modules representing T7SS in GBS from different ST’s (Figure 1).

**Table 1:** Type VII Secretion System components among different GBS isolates

|  |  | EsxA<br>SAG0230 | EsxB | SAG1032 | EssC | EssC | EssB | EsaB | EssA | EsaA | EsxA<br>SAG1039 |
| --- | --- | --- | --- | --- | --- | --- | --- | --- | --- | --- | --- |
| ST17 | a | 98.8 | 34.6 |  |  |  |  | 98.8 |  |  | 99.7 |
|  | b |  | 34.6 |  | 87.8 |  | 98.1 | 98.8 | 96.4 | 96.4 |  |
|  | c |  | 34.6 |  | 87.8 |  | 98.1 |  |  |  |  |
|  | d |  | 34.6 |  | 87.8 |  | 98.1 | 98.8 | 96.4 | 96.4 |  |
|  | e |  | 34.6 |  | 87.8 | 96.4 | 98.1 | 98.8 | 96.4 | 96.4 |  |
|  | f |  | 34.6 |  | 87.8 |  | 98.1 |  | 96.4 | 96.4 |  |
|  | g |  | 34.6 |  | 87.8 | 96.4 | 98.1 | 98.8 | 96.4 | 96.4 |  |
|  | h |  | 34.6 |  | 87.8 |  | 98.1 | 98.8 | 96.4 | 96.4 |  |
|  | i | 98.8 | 34.6 |  | 87.8 |  | 98.1 | 98.8 | 96.4 | 96.4 |  |
|  | j |  | 34.6 |  | 87.8 |  | 98.1 | 98.8 | 96.4 | 96.4 |  |
|  | k | 98.8 | 34.6 |  | 87.8 |  | 98.1 | 98.8 | 96.4 | 96.4 | 87.7 |
|  | l |  |  |  |  | 96.4 | 98.1 | 98.8 | 96.4 |  |  |
|  | m |  | 34.6 |  | 87.8 |  | 98.1 | 98.8 | 96.4 | 96.4 |  |
|  | n | 98.8 | 34.6 |  | 87.8 | 96.4 | 98.1 | 98.8 | 96.4 | 96.4 | 87.7 |
|  | o |  | 34.6 |  | 87.8 | 96.4 | 98.1 | 98.8 | 96.4 | 96.4 |  |
|  | p |  | 34.6 |  | 87.8 |  | 98.1 | 98.8 | 96.4 | 96.4 |  |
|  | q | 98.8 | 34.6 |  | 98.1 |  | 98.1 | 98.8 | 96.4 | 98.1 | 99.7 |
| ST1 | a | 98.8 | 34.6 | 61 | 97.2 |  | 98.1 | 98.8 | 96.4 | 98.1 | 99.7 |
|  | b | 98.8 | 34.6 | 61 | 97.2 |  | 98.1 | 98.8 | 96.4 | 98.1 | 99.7 |
|  | c | 98.8 | 34.6 | 61 | 97.2 |  | 98.1 | 98.8 | 96.4 | 98.1 | 99.7 |
|  | d | 98.8 | 34.6 | 61 | 97.2 |  | 98.1 | 98.8 | 96.4 | 98.1 | 99.7 |
|  | e | 87.8 | 34.6 | 61 | 97.2 |  | 98.1 | 98.8 | 96.4 | 96.4 | 99.7 |
|  | f | 98.8 | 34.6 | 61 | 97.2 |  | 98.1 | 98.8 | 96.4 | 98.1 | 99.7 |
|  | g | 98.8 | 34.6 | 61 | 97.2 |  | 98.1 | 98.8 | 96.4 | 98.1 | 99.7 |
|  | h | 98.8 | 34.6 | 61 | 97.2 |  | 98.1 | 98.8 | 96.4 | 98.1 | 99.7 |
|  | i | 98.8 | 34.6 | 61 | 97.2 |  | 98.1 | 98.8 | 96.4 | 98.1 | 99.7 |
|  | j | 98.8 | 34.6 | 61 | 97.2 |  | 98.1 | 98.8 | 96.4 | 98.1 | 99.7 |
|  | k | 98.8 | 34.6 | 61 | 97.2 |  | 98.1 | 98.8 | 96.4 | 98.1 | 96.4 |
| ST19 | a | 98.8 |  |  |  |  |  |  |  |  | 87.7 |
|  | b | 98.8 |  |  |  |  |  |  |  |  | 87.7 |
|  | c | 98.8 |  |  |  |  |  |  |  |  | 87.7 |
| ST6 | a |  |  |  |  |  |  |  |  |  |  |

**Figure 1.**
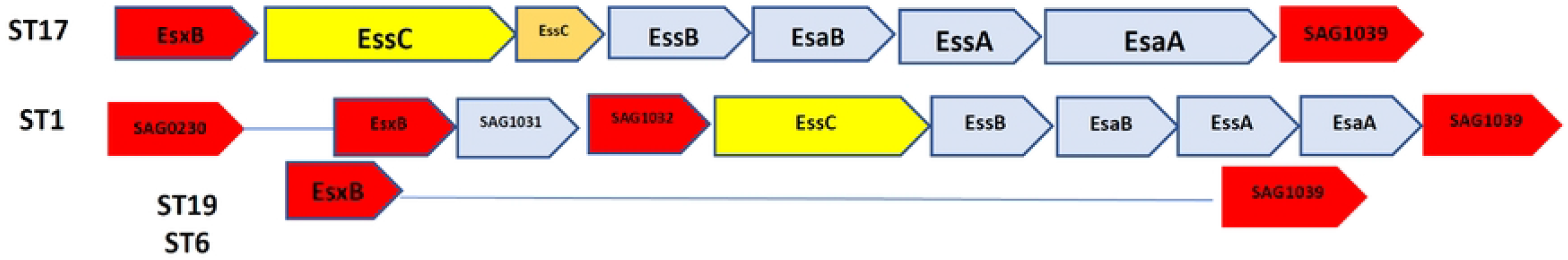
Comparison of the schematic structure of the T7SS system between different GBS ST types.

Protein percent identity of the T7SS components among various GBS STs. All ST17 were from neonates with EOD; ST1, ST 19 and ST6 GBS strains were obtained from colonized pregnant women. They were compared to the reference strain genome *S. agalactiae 2603 V/R*.

#### Module I (ST17, n=17)

ST17 isolates encoded two copies of the *essC* gene (SAG1003 and SAG1034); one copy of *esxA* encoding the WXG100 protein (present in only 23.5% of isolates) upstream of the T7SS core genes, and additional putative T7SS effector-*esxB* (including an LXG-domain containing protein) downstream of T7SS core genes.

#### Module II (ST1, n=11)

ST1 isolates encode one copy of *essC* gene (SAG1033); three copies of the WXG100 protein-encoding gene, *esxA*, SAG1039 located upstream, SAG1032 located downstream of the T7SS core genes, and another gene not directly linked to the T7SS locus that encoded a putative WXG100 protein (SAG0230). Like ST17 isolates, ST1 isolates encoded the *esxB* gene (T7SS effector including an LXG-domain containing protein) downstream of the T7SS core genes.

#### Module III (ST19, n=4; ST6 n=1)

all structural and regulatory T7SS genes were missing.

### Expression levels of essC and esxA genes among EOD and colonizing GBS isolates

We analyzed the transcription of genes encoding for the integral membrane bound ATPase protein *essC* (SAG1033) and one of the effectors, *esxA* (SAG1039), among EOD/ST17 (n=8) and colonizing/ST1 (n=8) GBS isolates. The *essC* gene was expressed in all tested GBS isolates (data not shown). However, *esxA* was weakly expressed among colonizing isolates compared to a significant expression in EOD isolates (data not shown).

#### GBS virulence (EOD and colonizing isolates) in *G. mellonella* model

We used *Galleria mellonella* larvae as an in vivo model of infection for GBS. The susceptibility of larvae to dose dependent killing by the WT GBS strain, *S. agalactiae* 2603 V/R (ATCC BAA-611), and two clinical EOD isolates was determined. We found that all isolates induced a dose-dependent response that was reproducible for each isolate in three independent experiments (data not shown).

We then injected varying doses of GBS isolates (EOD n=2, colonizing n=2) into each larva to compare the virulence of EOD and colonizing isolates by measuring the infecting dose (LD_50_). LD_50_ values obtained for infection with EOD isolates (2.7×10^6^) were significantly lower (p<0.05) than those of colonizing (ST1) isolates (4.1×10^8^), indicating that an isolate associated with EOD has increased virulence in *G. mellonella* compared to colonizing strains. Twenty-four hours after infection with EOD isolates, only 50% of infected larvae survived, compared to 85% survival rate after infection with colonizing strains (Figure 2).

**Figure 2.**
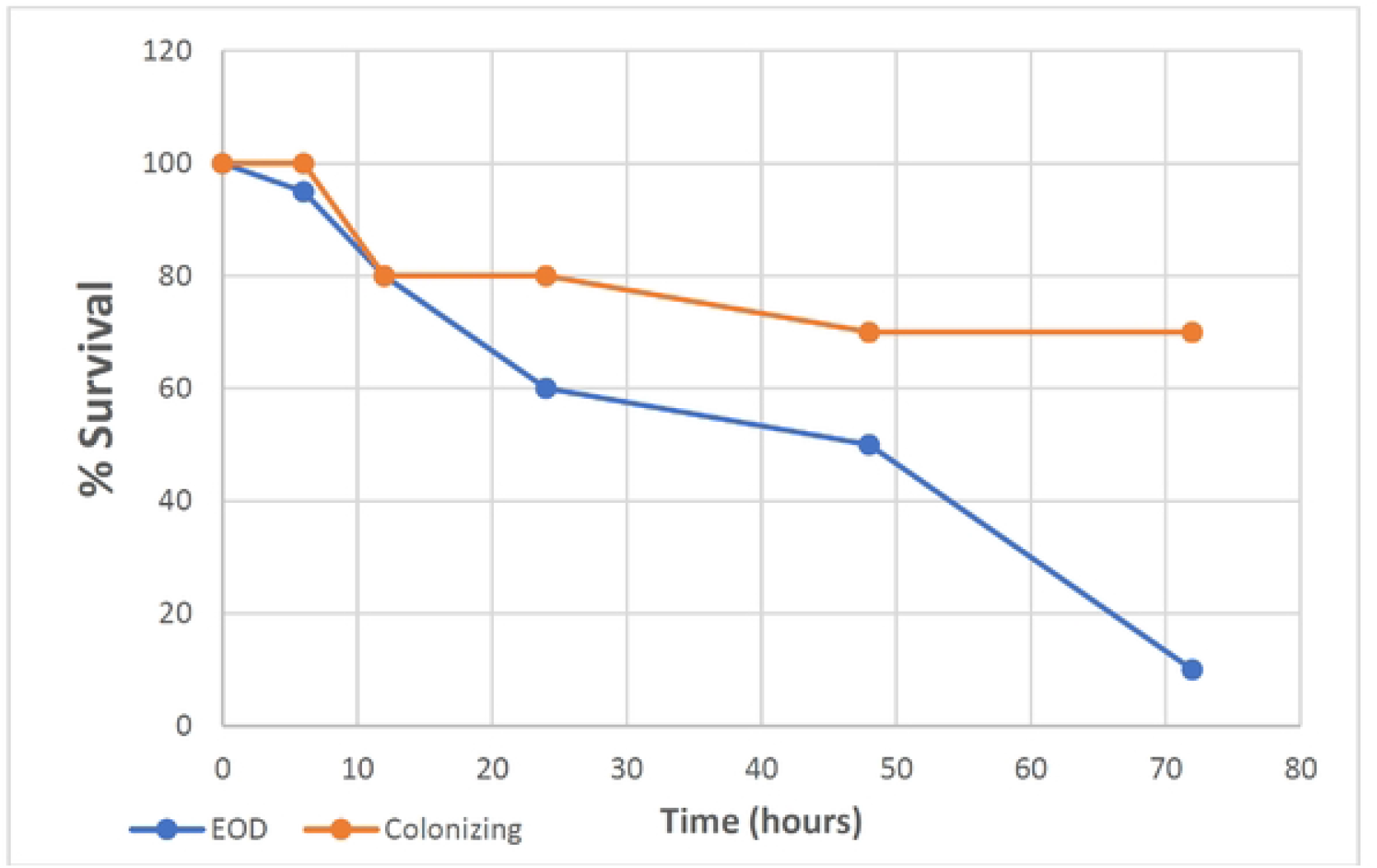
Kaplan-Meier survival curves of larvae challenged with EOD and colonizing isolates. A Kaplan–Meier survival plot of survival after infection with either EOD or colonizing strain. Data were collected from eight distinct experiments (four experiments with EOD strains and four experiments with colonizing strains) with 10 larvae per group for each experiment. Survival curves show one representative experiment, with use of 10 larvae per group. PBS-injected larvae were used as a negative control, and all survived until the endpoint of the experiment.

### Attenuation of 118659 EOD/ST17 isolate by essC knockout

To understand the role of EssC in virulence of GBS strains, we generated an isogenic *essC* mutant in the clinical isolate 118659 EOD/ST17. PCR analysis of the mutant 118659Δ*essC* produced bands with a different size than those observed for the 118659 WT strain (2900 bp versus 3300 bp), indicating that the *essC* gene was disrupted by the insertional mutagenesis of the kanamycin cassette (Figure 3a). The insertion of kanamycin resistance gene was validated using primer pairs v-omega-Km1 and v-omega-Km2, in composition with EssC-KO primers located in both ends of original amplicon. The PCR analysis using these primers generated bands only in the mutant strain and were absent in the WT strain (Figure 3b). According to variant analysis of the 118659 WT and 118659Δ*essC* (mutant) genomes against the reference genome, the difference between them relied only on the deletion of the *essC gene* and no additional mutations were identified. In standard rich medium the Δ*essC* had a similar growth rate as the WT 118659 (Figure 4). Additionally, Δ*essC* did not show a growth defect or difference on hemolysis production when cultured in parallel with the WT strain. All tested strains had the same prototypical phenotype and displayed a narrow zone of beta-hemolysis on blood agar plate.

**Figure 3.**
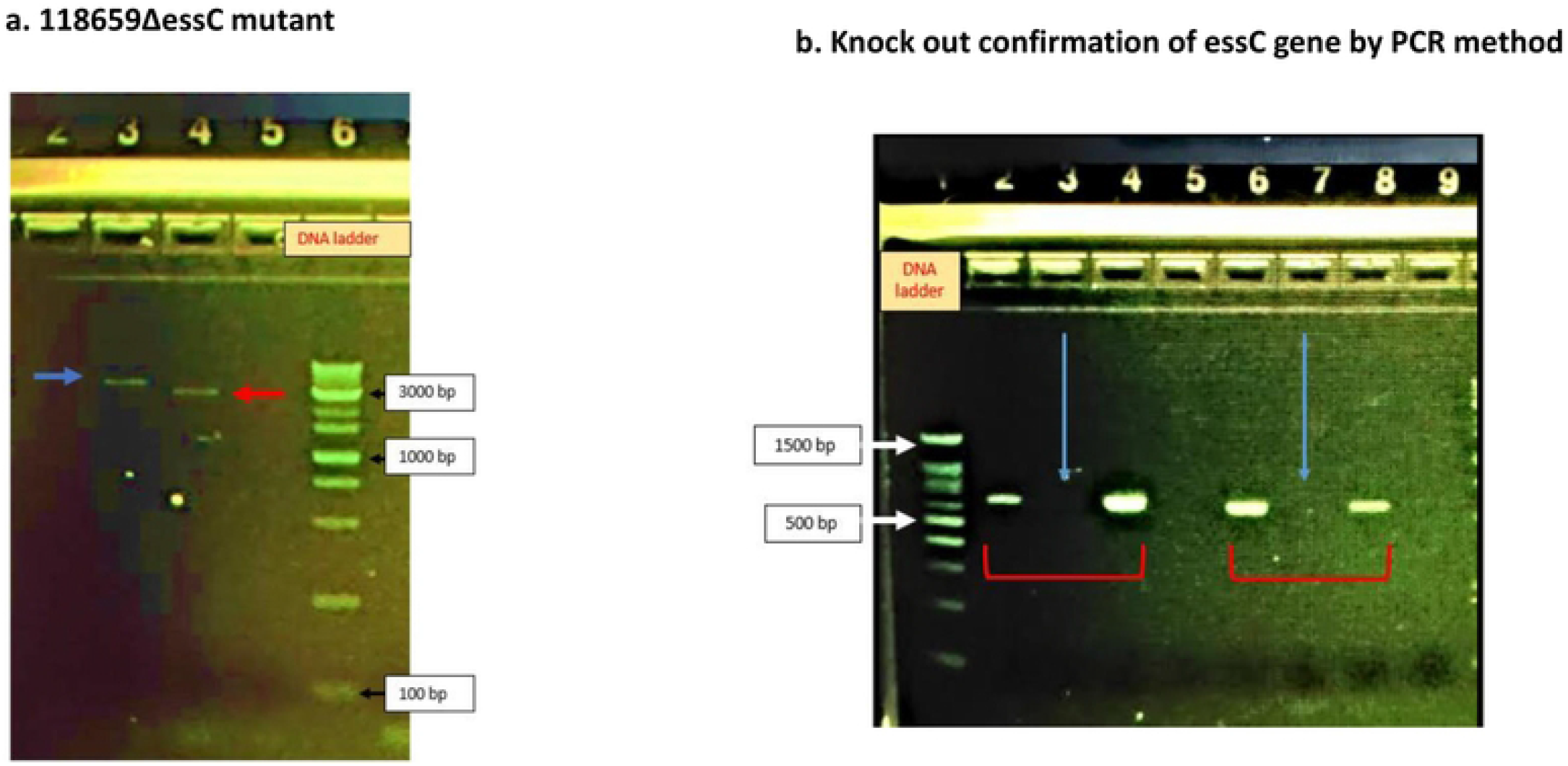
118659*ΔessC* mutant knockout. a. Knock out confirmation of *essC* gene by PCR method, using EssC-KO -F and EssC-KO -R primers flanking the *essC* gene. Lane 3 −118659 WT strain (~ 3300 bp.); Lane 4 - 118659*ΔessC* (2900 bp.); Lane 6 – 1 kb DNA Ladder. b. Knock out confirmation of *essC* gene by PCR method. Lane 1–100 bp DNA adder; Lane (2,4): 118659*ΔessC* mutant, with primers EssC-KO-F flanking the *essC* gene and v-omegaKm1 flanking the kanamycin resistance cassette (~ 621 bp.); Lane (6,8): 118659*ΔessC* strain, with primers EssC-KO-R flanking the *essC* gene and v-omegaKm2 flanking the kanamycin resistance cassette (~ 574 bp.). For comparison, the absence of band was identified with 118659 WT strain (Lane 3 and 7).

**Figure 4.**
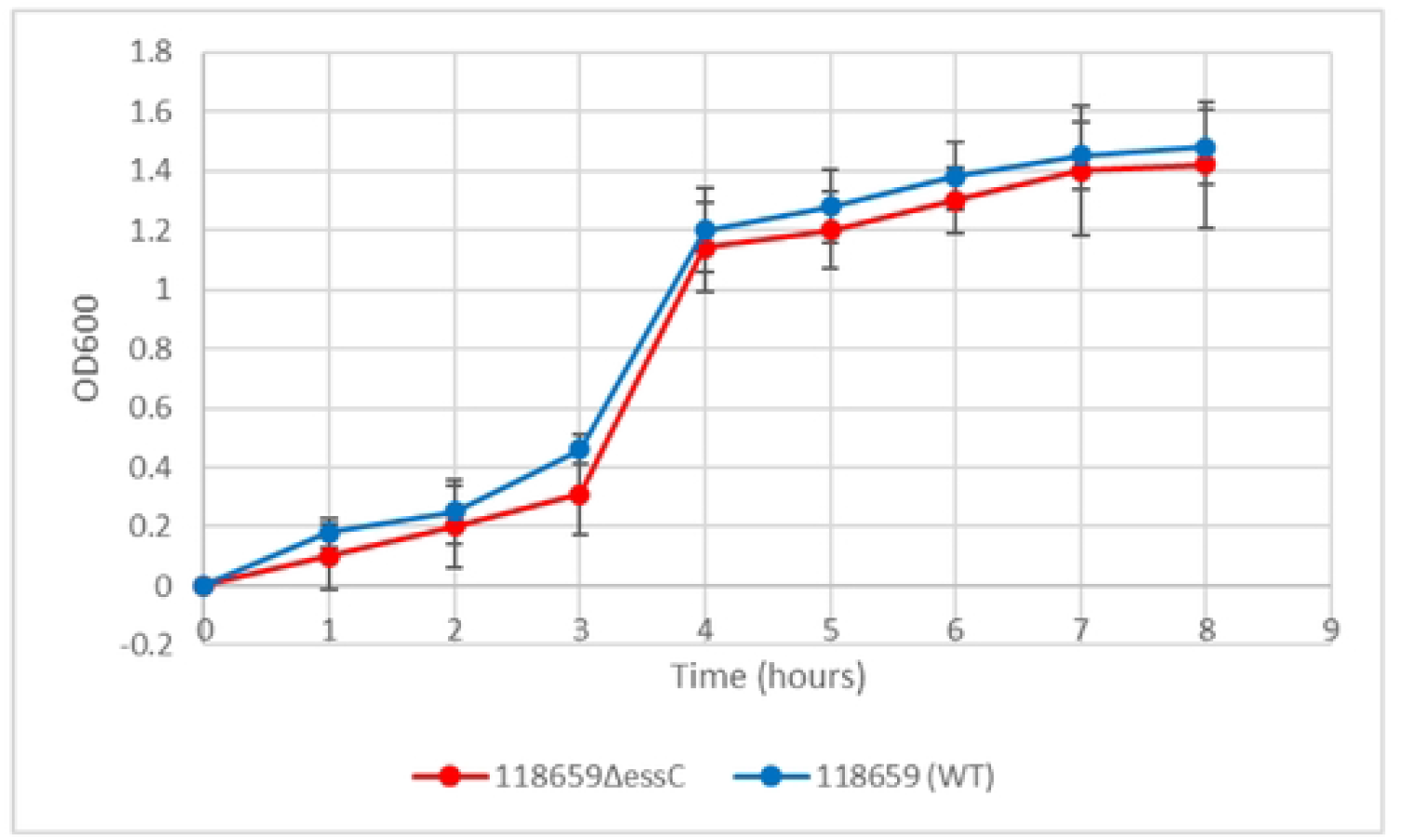
Growth rate of the wild-type 118659 (WT) and the mutant 118659*ΔessC* isolates Growth rate of the wild-type 118659 (WT) (blue line) and mutamt 118659*ΔessC* (red line) strains in BHI medium for 8 hours. Assays were repeated three times and are presented as mean ± SD. EOD strains showed increased virulence in *G.mellonella*. LD_50_ values determined by Probit analysis following infection of larvae by (A) EOD strains (blue) and colonizing strains (red). Each datapoint represents the LD_50_ of each experiment in which groups of 10 larvae were infected with four different inoculums.

### Expression of core components of T7SS in 118659ΔessC (mutant) strain

We compared the gene expression of *esaA, essA, essB*, and *esaB*, located upstream to *essC* gene to study the influence of essC knock out on their activity. qRT-PCR analysis revealed similar levels of expression of tested genes among mutant and WT strains demonstrating that the activity of whole T7SS locus was not disturbed by knocking out the *essC* gene.

### Contribution of essC gene mutation to GBS virulence in G. mellonella in vivo model

To assess the ability of the *G. mellonella* model to discern changes in virulence beyond the 118659Δ*essC* (mutant) and 118659 (WT) strain the infecting dose (LD_50_) for each strain was determined. LD_50_ values obtained for infection with mutant strain were significantly higher than those of WT strain (4.1×10^9^ compared to 2.7×10^7^, p < 0.01) indicating that in the *G. melonella* model the mutant strain is less virulent. Larval mortality appeared 6-8 hours after infection with both WT and the mutant isolates but increased progressively mainly in the WT strain. Larval mortality in the mutant inoculatec group was significantly reduced (p = 0.03) compared to the WT strain (Figure 5). The larval survival rate with WT strain infection was 10%, compared to 40% with the mutant strain. In summary, the mutant strain has decreased ability to kill *G. mellonella*, indicating that the *essC* gene may play an essential role in GBS virulence.

**Figure 5.**
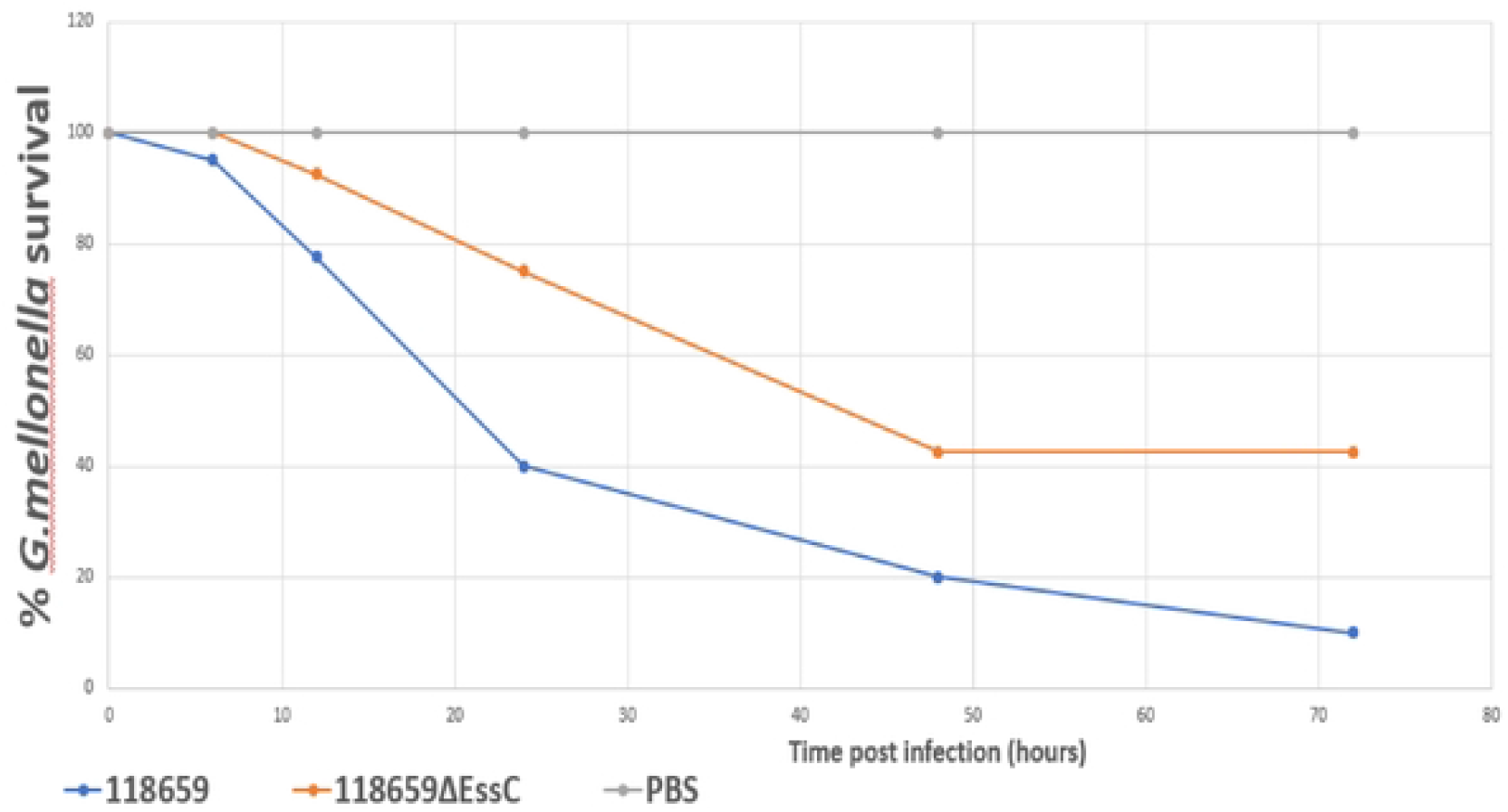
Kaplan-Meier survival curves of larvae challenged with 118659 (WT) and mutant strain (118659ΔEssC) Kaplan-Meier survival curves of larvae challenged with an inoculum of 10^7^ CFU of 118659 (WT) and mutant strain (118659ΔEssC), and PBS (control). Each infection was repeated three times with 10 larvae for each experiment, (p < 0.05; log-rank test).

### Kinetics of in vivo growth of WT and GBSΔessC (mutant) strains

To monitor growth of GBS in infected larvae, the groups of 10 larvae were infected with 118659 (WT) and 118659ΔessC (mutant) strains (~1×10^6^ or ~1×10^8^ CFU/larvae, respectively), and bacterial burden was measured hourly in pools of larvae. During the first 12 hours, the larval burden of both WT and mutant isolates increased over time and reached to ~1 × 10^10^ CFU (Figure 6). After 12 hours, the burden of the WT strain decreased faster compared to the mutant strain. Larvae that outlived the infection with the mutant strain over 24 hours seemed to clear the GBS [10^7^ CFU/mL 24 h p.i.; 10^6^ CFU/mL 48 h p.i.]. This is probably due to efficient phagocytosis of larval hemocytes [121,122]. Finally, after 72 hours, the larval burden in the mutant strain was ≈ 3 logs higher compared to the WT strain (10^3^ CFU/mL to 10^1^ CFU/mL).

### G. mellonella health index following infection with 118659ΔessC strain

To measure more subtle differences in larvae health status post-infection with 118659 (WT) and 118659Δ*essC* (mutant) strains, *G*. *mellonella* larvae were monitored daily for the following attributes: activity, extent of silk production (cocoon formation) and melanization (Table S4). Higher activity and increased cocoon formation corresponded to healthier larva. In our experiments, the activity of the *G. mellonella* larvae was similar for both strains WT and mutant isolates. Melanin production occurred as fast as 6 hours after infection with the mutant strain and proceeded until the end of the experiments (72 hours) (Table S5). Melanin production was not fully correlated with mortality of the larvae. We found live larvae with full melanization even after 72 hours. Larvae infected with mutant GBS strain were able to produce more cocoon compared to larvae infected with the WT strain, even when the melanization process already started. Healthy larva received a score of 7-8 points, while very sick larvae received a low score (<5). WT strains caused increased melanization, lower activity and cocoon formation, and were associated with a low health index – score 0 (72 h after inoculation) of *G. mellonella*. In contrast, the mutant strain caused an intermediate infectious process (72 hours after inoculation) with a health score of approximately 2.

Thus, larvae infected with mutant GBS strains received higher health scores. These larvae successfully produced cocoons, even during progressive melanization and overcame the infection.

### Decreased fitness of mutant in the G. mellonella model

To determine relative differences in strain fitness of 118659Δ*essC* (mutant) compared to 118659 (WT) GBS strains, we performed a competition assay using *G. mellonella*, which could be more sensitive in detecting changes in bacterial fitness, than the survival assay^32^. To distinguish between WT and mutant GBS strain, we induced resistance to streptomycin (Sm) in the WT strain by culturing and passaging it several times under high streptomycin concentrations. The mutant strain showed decreased fitness in *G. mellonela* model with 36.2% (SD 2.215) of the recovered CFU belonging to the mutant strains compared to 63.8% of WT (SD 2.375), p < 0.0001. As a control, to make sure that homogenization did not impact relative bacterial survival, we plated a portion of the initial mixed culture prior to injection into the larva and saw no difference in relative survival between the wild-type and the mutant strains (data not shown). According to our results, there are relative differences in strain fitness of WT and mutant strain, which could confirm the decreasing virulence of the mutant strain.

### Bacterial clearance by G. mellonella

To study the difference in bacterial clearance by *G. mellonella* after infection with sublethal doses of the mutant and WT strains larvae were injected with a sublethal inoculum ≈1 × 10^5^ CFU of 118659 (WT) strain and ≈1 × 10^6^ CFU 118659Δ*essC* of the mutant strain. The larvae were monitored every hour for 7 hours and after 12 hours. During the first 8 hours post infection, the bacterial burden of the WT in the larvae rapidly increased to three logs compared to the initial inoculum (Figure S1), but then decreased back to the initial levels. In contrast, the bacterial burden of the mutant strain in the larvae failed to multiply in the same rate and the eventually, the bacterial burden increased by only one log. Overall, the bacterial burden of the mutant strain was relatively stable over time. To conclude, we demonstrated by the competition assay differences in bacteria fitness of the WT and mutant isolates.

## DISCUSSION

In this study we performed a genomic survey of T7SS in clinical GBS isolates, obtained from blood cultures of neonates with EOD and collected from vaginal screening of asymptomatic pregnant women. T7SS has been well characterized in *Mycobacterium* species, in terms of its structure, functions, and transport models^11^. Recent advances have also facilitated our understanding of T7SS in GBS^26,27^. Here, we compared the structure of T7SS locus among EOD/ST17 and colonizing/ST1 GBS isolates. We identified significant differences in the structure of T7SS between EOD and colonizing GBS strains. Notably in 76.5%% of EOD/ST17 strains putative effectors were abscent: *esxA* (WXG100 protein-encoding gene) and *esxB* (LXG-domain containing protein), while in colonizing/ST1 isolates three copies of the WXG100 protein-encoding gene *(esxA*) and one copy of the essC gene (SAG1033) were observed. In contrast to the type-specific capsular polysaccharides which are well-defined virulence determinants^4^, the role of WXG100 proteins and LXG-domain containing protein as a virulence factor is not yet clearly understood. These proteins may enhance the human immune response to GBS infection. Absence of *esxA* and *esxB* genes in most ST17 isolates, may protect them from opsonization and killing by humoral and cell-mediated processes in the host. In several colonizing isolates (ST6 and ST19) structural and regulatory genes encoded by T7SS locus were missing. The ST-dependent T7SS diversity in GBS was recently described by Zhou et al^27^.

In our study we highlight the diversity of T7SS in relation to clinical syndromes. We compared the virulence of EOD and colonizing isolates using *G. mellonella* larvae, an in vivo model of GBS infection. The use of *G. mellonella* larvae as bacterial infection model was developed as an alternative to murine or other vertebrate infection models to contribute to the 3Rs (reduction, replacement, and refinement) of animal use in scientific research^38^. In vivo larval experiments demonstrated a difference in the pathogenicity of various clinical GBS strains. GBS strains associated with EOD demonstrated enhanced virulence in *G. mellonella* compared to colonizing strains. These results are consistent with previous study, where GBS disease associated isolates were able to establish systemic infection of G. *mellonella* larvae with extensive bacterial replication and dose-dependent larval survival^32^.

We further demonstrated the role of T7SS in virulence of ST17 strains and showed that it dependends on the proper activity of EssC, a membrane-embedded ATPase of the FtsK/SpoIIIE family. We generated an EOD/ST17 mutans by knocking out the *essC* gene and compared the virulence of the mutant and WT strains in *G. melonella* larvae in vivo model. According to our results, the knocked-out mutant 118659*ΔessC* has reduced ability to kill *G. mellonella*. Furthermore, LD_50_ values obtained with the 118659*ΔessC* strain were significantly higher than those obtained with the WT 118659 strain. Our results are in line with a recently published study^26^, which demonstrated that deletion of the ATPase-encoding gene, *essC*, mitigates virulence and GBS-induced inflammation in the brain, as well as cell death in brain endothelial cells in murine model of hematogenous meningitis.

Consistent with this, our data indicates that *EssC* deletion affected bacterial growth during infection, as well as bacterial fitness and the response of larvae to GBS infection. We show that larvae were more effective in eradicating 118659*ΔessC* strain infection and this is probably related to different immune responses^39^. The competition model for *G. mellonella* was found as more sensitive in discerning relative differences in *Bacillus anthracis* strain fitness than the survival assay^36^. In our competition assay, the mutant strain showed decreased fitness, which could confirm the decreasing virulence of the mutant strain. The possible explanation is that T7SS in GBS secrete various effectors which induce immune tolerance against GBS infection. In the mutant strain (118659Δ*essC*) lacking the functional secretion system, the larvae’s immune system is more effective in eradication of the mutant strain^39^. Finally, we show that *EssC* deletion was associated with an increase in the health index of *G. mellonela* during infection, regarding activity, cocoon formation and melaninization. The health index scoring system evaluates the health status of the larva during an infectious process. This parameter is also used to measure differences in virulence of other bacterial pathogens in *G. mellonella^40,41^*. We show that melanin production by the larvae infected with the mutant strain occurred very quickly (after 6 hours p.i.). Although melaninization is usually associated with imminent death of the larva, larva remained viable, and even succeeded to produce a cocoon. We think this indicates that the mutant strain cannot succeed in causing massive dissemination of infection. This indicates that the mutant strain is attenuated compared to the WT as other parameters such as the larva’s immune function and the infective dose were similar between experiments.

In conclusion, our findings indicate that the T7SS plays an essential role during infection and contributes to GBS pathogenicity. The proper function of T7SS, by efficient secretion of various effectors could be considered as a virulence factor of invasive GBS isolates. In most of our ST17 isolates the genes encoding to classical T7SS effector *(esxA, esxB*) were absent. This may be related to their ability to escape from the immune system. Our results establish a link between T7SS and EOD in the newborn and may partially explain, why in most colonized women colonization does not proceed to infection in the newborn. Further studies are warranted to identify other effectors, their effect on substrate recognition and specificity, the inflammasome and immune response.

## MATERIALS AND METHODS

### Bacterial strains and growth conditions

A total of 33 GBS clinical isolates obtained from blood cultures of neonates with EOD (n=17) and GBS isolates collected from the vagina of asymptomatic pregnant women (n=16) were studied (Table S1). GBS strains were grown in BHI medium (Hylabs, Israel) at 37°C with 5% CO2 under shaking conditions. *Escherichia coli* was grown aerobically in Luria–Bertani (LB) (Hylabs, Israel) at 37°C. Antibiotics were added: for GBS 250 μg/ml kanamycin (Km), and 1 μg/ml erythromycin (Em); for *E. coli:* 100 μg/ml ampicillin (Amp), 500 μg/ u ml Em and 50 μg/ml Km. All antibiotics were purchased from Sigma-Aldrich (St Louis, MO, USA).

### Bioinformatic analysis of T7SS genes in GBS clinical isolates

Genomic libraries of clinical GBS isolates were prepared using Nextera XT kits (Illumina, San Diego, CA) and sequenced using the Illumina MiSeq Reagent Kit v3 (600-cycle). The reads obtained for each sample were trimmed and the quality of the Fastq reads was examined using the Fastq Utilities Service, and finally assembled by SPAdes using the PATRIC website^29^. The presence of T7SS genes was identified using web-resources: the bacterial bioinformatics database and analysis resource of PATRIC website (https://www.patricbrc.org/)and NCBI BLASTp (available at www.ncbi.nlm.nih.gov/blast/). A high-quality representative genome of *Streptococcus agalactiae* 2603V/R ATCC BAA611 (serotype V, ST19) was used as reference^30^. We characterized the structure and membrane topology of genes using the HHpred interactive server. We identified genes that encode WXG100 proteins, that are presumably secreted by T7SS, by detection the presence of signal peptides using Phobius and SignalP tools. We compared the structure and the presence of T7SS effectors among ST17 and ST1 GBS isolates.

### Generation of knockout GBS strain

Deletion mutant was created using the temperature sensitive plasmid pJRS233 with a kanamycin resistance gene, Km in the knockout construct, as previously described^31^. Briefly, the flanking region of *essC* gene of GBS118659 were amplified using EssC-KO-F and EssC-KO-R primer pairs (Table S2). The 3809-bp PCR product was purified and cloned into pGEM-T-Easy (Promega, Medison WI, USA) to yield pGEM: *essC*. The plasmid was transformed to *E. coli* DH5α by electroporation, plated on LB plates containing ampicillin 100 μg/mL with x-gal and IPTG, and allowed to grow for one day at 37°C. Positive transformants (white colonies) were confirmed by PCR and sequencing. Restriction of pGEM: *essC* plasmid with HpaI and KpnI, releases a 2934 fragment of *essC* leaving 411bp and 465bp of *essC* on each side-for homologous recombination to GBS chromosome. Next, the digested plasmid was treated with Klenow enzyme and ligated with a 2043bp fragment of *SmaI* digested Ωkm cassette (kanamycin resistance cassette flanked by Ω elements). The pGΔEssCΩKm plasmid was transformed to *E. coli* DH5α by electroporation, plated on LB plates containing kanamycin (Km)-50 μg/mL. Positive transformants were confirmed by PCR and sequencing. The plasmid was restricted with NotI (releasing a 2954 bp fragment of ΔEssCΩKm flanked by essC sequences) and ligated into NotI digested pJRS233 plasmid (a temperature-sensitive shuttle vector). The pJΔ: EssC: ΩKm plasmid was transformed to *E. coli* DH5α by electroporation, plated on LB plates containing Km50/Em500. Positive transformants were confirmed by PCR. 3-7μg of pJΔ: EssC: ΩKm plasmid was transformed into competent GBS cells (strain 118659) by electroporation (25 μF, 400ohms, 1.75 KV) and bacteria were plated on THY plates containing erythromycin 1 μg/mL. Erythromycin-resistant transconjugants were then cultured under non-permissive temperature to select for single cross-over recombinants, followed by serial passage in antibiotic-free BHI and screening for double cross-over deletion mutants by PCR. Deletion was confirmed by PCR amplification of the regions spanning the deleted fragment using the EssC-KO-F and EssC-KO-R primers, primers for kanamycin resistance, and pair of primers from inner part of *essC* gene (Conf-KO-essC), which should be replaced by omega kanamycin cassette (Table S2). The absence of any secondary site mutations was confirmed by whole genome sequencing.

### In vitro phenotypes of 118659 ΔessC mutant and 118659 Wild type (WT) strains

The 118659 Wild type (WT) and 118659Δ*essC* (mutant) strains were grown overnight in BHI medium, 1:20 diluted in fresh BHI medium at the zero-time point and incubated at 37°C + 5% CO2 under shaking conditions. The optical density (OD) at wavelength 600 nm of each group, was measured for 8 hours (achieving the stationary phase). Each experiment was repeated three times. The WT and mutant strains were cultured on blood agar (Hylabs, Israel) and incubated at 37°C + 5% CO2 for 24 hours to observe hemolytic activity.

### Galleria mellonella in vivo model

*G. mellonella* larvae were obtained from Volcani center (Dr. Dana Ment laboratory, Entomology department), kept in darkness at room temperature, and discarded after one week following arrival. Healthy larvae measuring from 2-2.5 cm were used for all experiments. Injections were done using INSUMED 29G insulin syringes (Pic solution)^32^. For each experiment groups of 10 larvae were injected with 10 μl of serial dilutions of bacterial suspension. A control group including five larvae were inoculated with PBS for control of motility change caused by physical injury or infection by a contaminant. Experiments were repeated twice. After injection, larvae were observed at room temperature for 15–30 min to ensure recovery and were stored in Petri dishes in the dark at 37 °C. Survival of infected larvae was monitored for 72 hours post infection (p.i). The larvae were considered dead when non-responsive to touch.

### Survival assay

GBS isolates were grown to an OD 0.4-0.6 in BHI (~1 × 10^9^ colony forming units [cfu] per ml), washed and resuspended in PBS (Hylabs, Israel), and then diluted prior to injection. Cells were washed twice in sterile PBS and diluted to the desired inoculum. The starting inoculum was confirmed through serial dilution, plating on blood agar plates (Hylabs, Israel) just before administration for CFU counting. For the determination of the infecting dose (LD_50_), four groups of 10 larvae were injected with 20 μl of serial dilutions of bacterial suspension as described above. Survival curves were plotted using Kaplan–Meier method and differences in survival were calculated using the log-rank test (SPSS). LD_50_ was calculated using the Probit method and differences in LD_50_ between different isolates were assessed using the Mann-Whitney test.

### In vivo GBS growth curve

Groups of 10 larvae were infected with 118659 (WT) and 118659*ΔessC* (mutant) strains and monitored for 72 hours. At fixed time points (8, 24, 48, and 72 h p.i.), larvae were kept at −20°C for 10 min before being transferred to Eppendorf containing 100 μL of sterile PBS, homogenized by mechanical disruption, serially diluted. CFU counts from homogenized infected larvae were determined by viable plate count method using selective Chromo Strep B plates (Hylabs, Israel).

### Competition assay

To distinguish between WT and mutant GBS strain, we induced resistance to streptomycin (Sm) in the WT strain by culturing and passing it several times under high streptomycin concentrations. The GBS strains were grown to log phase (OD_600_= 0.4) for 3-4 hours, washed and resuspended in PBS. Mutant strains were mixed with the parental (WT) at a 1:1 ratio. Ten microliters of the mixed culture (~1 × 10^7^ total CFU) were injected into each larva and larvae were then incubated for 24 h at 37°C. We chose 24 hours as this was long enough for the infection to become established but short enough to preclude total larval mortality. The larvae were then rinsed in 70% ethanol followed by sterile water to help minimize contamination by surface bacteria before being homogenized in PBS by mechanical disruption. Homogenates were plated on BHI and BHI-antibiotic plates (Hylabs, Israel) (streptomycin (SM500) for WT and kanamycin (Kan250) for mutant strain) and the CFU recovered for each strain was calculated.

### Monitoring of G. mellonella larvae

Each *G*. *mellonella* larvae were monitored daily for activity, silk production (cocoon formation) and melanization (Table S4). Loh et al ^33^ developed these criteria to evaluate the health status of the larva during an infectious process. This parameter is used to measure more subtle differences in virulence of different bacterial pathogens in *G. mellonella* ^33–36^. An uninfected group and a group inoculated with saline were used as negative controls. A score was assigned to each observation, and an overall health index score was calculated for each larva.

### Clearance of mutant and WT strains by G. mellonella

*G. mellonella* larvae were injected with a sublethal inoculum (the closest dose to killing 15% of the larvae) ≈1 × 10^5^ CFU of 118659 (WT) strain and ≈1 × 10^6^ CFU 118659Δ*essC* (mutant) strain, monitored every hour for 7 hours and after 12 hours. At each fixed time point, three surviving larvae were randomly selected, kept for 15 min on ice and bathed in 70% ethanol and sterile water. The selected larvae were homogenized in 2 ml. For bacterial count serial dilution were performed and the homogenate was plated in blood agar (Hy-Labs, Israel) and selective Chromo Strep B plates (Hy-Labs, Israel).

### Transcriptional analyses

Quantitative RT-PCR analysis of *esxA*, *essA*, *essB*, *esaB* and *essC* genes expression was performed as described previously^27^. Primers were designed using Primer3 Plus and Clone manager 9 professional edition, ver 9.4 software. Primers were used at a final concentration of 0.4 μmol/L (Table S3). RNA was extracted from GBS cultures grown at 37°C to an exponential growth phase in BHI medium. RNA was purified using the Rneasy Mini kit (Qiagen) according to manufacturer instructions. Purified RNA was treated with the DNAse kit (HY-labs, Israel) according to manufacturer instructions. The RNA quality and concentration was assessed by NanodropTM and visually on a 2 % E-Gel with SYBR safe (Invitrogen, Thermo) and visualized by E-Gel Power Snap Electrophoresis device (Invitrogen, Thermo Fisher). cDNA was synthesized using the Hy-RT-PCR kit (HY-labs, Israel), according to manufacturer instructions. cDNA was diluted 1:150 to further reduce bacterial DNA contamination and qPCR was performed using Hy-SYBR power mix (HY-labs, Israel) and CFX96 Real-Time System (Biorad). RNA from three independent biological triplicates were analyzed and final cycle threshold for each strain was calculated (mean value of three experiments). Relative quantification of gene expression was performed using comparative 2^-ΔΔ^Cτ. Results were normalized using *rpoB* gene as the housekeeping gene.

### Statistical analyses

Statistical analysis was performed using SPSS version 27.0 (SPSS Inc., Chicago, IL, USA). Statistical details of experiments, such as statistical test used, experimental *n*, can be found in each figure legend. Significance was defined as p < 0.05.

## ACKNOWLEDGEMENTS

The authors report there are no competing interests to declare. The study was funded by internal funds of the Microbiology laboratory, Mayaney Hayeshua, Bney Brak, Israel and the Infectious Disease Unit, Sheba Medical Center.

## Legends to figures

**Figure S1.**
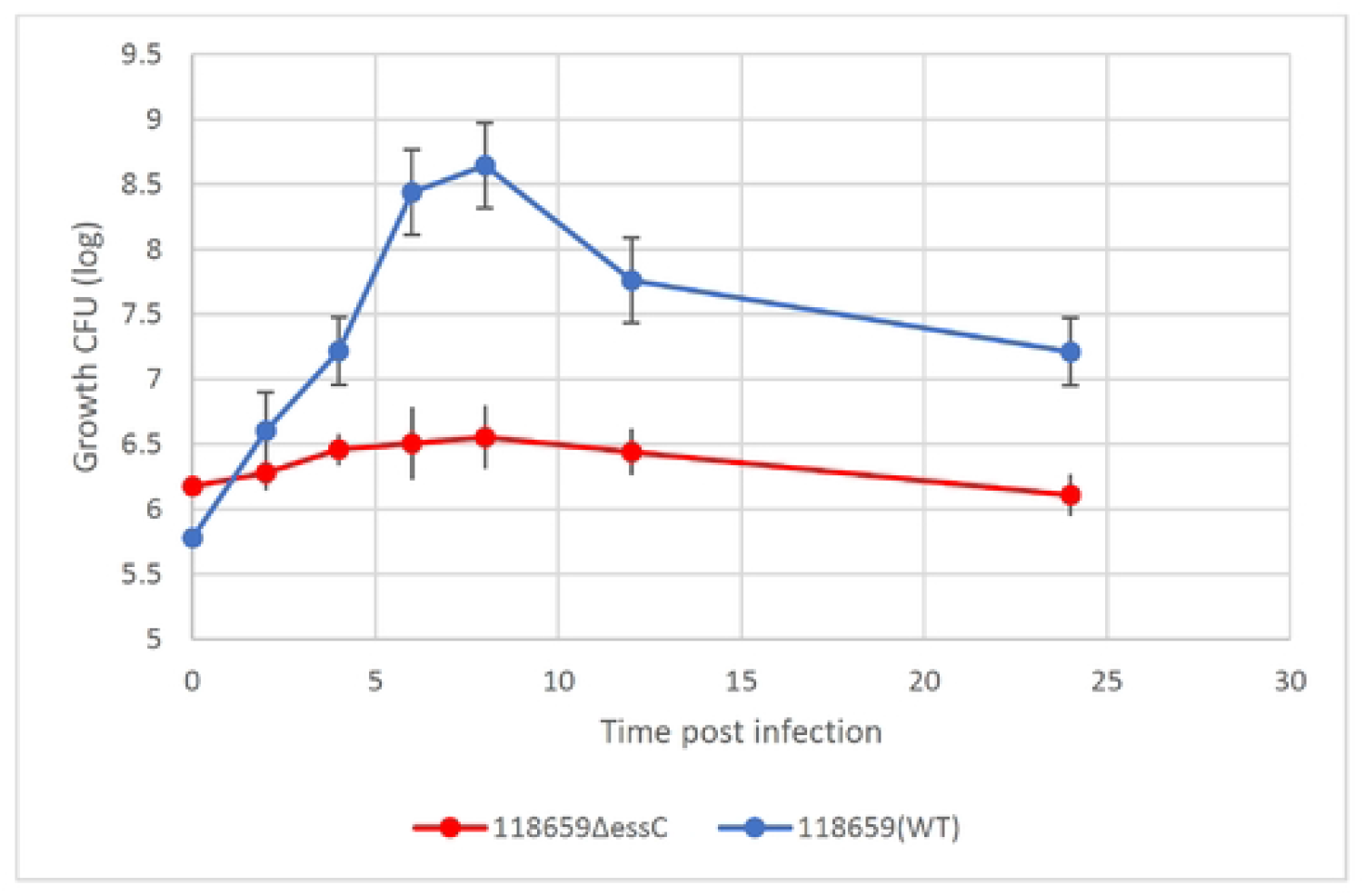
Kinetics of 118659 (WT) and H8659ΔEssC bacterial growth in vivo

